# DivIVA controls the dynamics of septum splitting and cell elongation in *Streptococcus pneumoniae*

**DOI:** 10.1101/2024.05.09.593393

**Authors:** Jennyfer Trouve, André Zapun, Laure Bellard, Dimitri Juillot, Anais Pelletier, Celine Freton, Morgane Baudoin, Rut Carballido-Lopez, Nathalie Campo, Yung-Sing Wong, Christophe Grangeasse, Cecile Morlot

## Abstract

Bacterial shape and division rely on the dynamics of cell wall assembly, which involves regulated synthesis and cleavage of the peptidoglycan. In ovococci, these processes are coordinated in an annular mid-cell region with nanometric dimensions. More precisely, the cross-wall that is synthesized by the divisome is split to generate lateral wall, whose expansion is insured by insertion of so-called peripheral peptidoglycan by the elongasome. Septum cleavage and peripheral peptidoglycan synthesis are thus crucial remodeling events for ovococcal cell division and elongation. The structural DivIVA protein has long been known as a major regulator of these processes but its mode of action remains unknown. Here, we integrate click chemistry-based peptidoglycan labeling, direct stochastic optical reconstruction microscopy and in silico modeling, as well as epifluorescence and stimulated emission depletion microscopy to investigate the role of DivIVA in *Streptococcus pneumoniae* cell morphogenesis. Our work reveals two distinct phases of peptidoglycan remodeling along the cell cycle, that are differentially controlled by DivIVA. In particular, we show that DivIVA ensures homogeneous septum cleavage and peripheral peptidoglycan synthesis around the division site, and their maintenance throughout the cell cycle. Our data additionally suggest that DivIVA impacts the contribution of the elongasome and class A PBPs to cell elongation. We also report the position of DivIVA on either side of the septum, consistent with its known affinity for negatively curved membranes. Finally, we take the opportunity provided by these new observations to propose hypotheses for the mechanism of action of this key morphogenetic protein.

**Importance:** This study sheds light on fundamental processes governing bacterial cell growth and division, using integrated click chemistry, advanced microscopy and computational modeling approaches. More precisely, it addresses mechanisms involved in the regulation of cell wall synthesis and remodeling in *Streptococcus pneumoniae*. This bacterium belongs to the morphological group of ovococci, which includes many human pathogens, such as streptococci and enterococci. In this study, we have dissected the function of DivIVA, which is a structural protein involved in cell division, cell morphogenesis and chromosome partitioning in Gram-positive bacteria. This work unveils the role of DivIVA in the orchestration of cell division and elongation along the pneumococcal cell cycle. It not only helps understanding how ovoid bacteria proliferate, but also offers an opportunity to consider how DivIVA might serve as a scaffold and sensor for particular membrane regions, and thus be involved in various processes associated with the cell cycle.

## Introduction

Peptidoglycan (PG) is a three-dimensional network made of repeated *N*-acetylglucosamine-*N*-acetylmuramic acid disaccharide units cross-linked by short peptides (Vollmer et al., 2008a). PG confers resistance to turgor pressure and a specific cellular shape adapted to the environment of the bacterium. This shape is tuned by the dynamics of PG synthesis and remodeling (Pinho et al., 2013; Trouve et al., 2021b), whose enzymatic actors are well known but whose organizational and regulatory factors remain poorly understood.

Assembly of the PG at the cell surface is performed by transglycosylase activities, which polymerize the PG precursor unit lipid II (a membrane-anchored disaccharide pentapeptide) into glycan chains, and by transpeptidase activities, which crosslink the peptide chains. PG synthases include SEDS proteins (Shape, Elongation, Division and Sporulation), which carry transglycosylase activities, and Penicillin-Binding Proteins (PBPs) (Meeske et al., 2016; Sauvage et al., 2008). Class A PBPs (called aPBPs henceforth) are bifunctional enzymes carrying both transglycosylase and transpeptidase domains, while class B PBPs (bPBPs) are monofunctional transpeptidases (Meeske et al., 2016; Sauvage et al., 2008). bPBPs work in tandem with a cognate SEDS protein and participate to the cell division or cell elongation complexes called the divisome or the elongasome, respectively (Käshammer et al., 2023; Reichmann et al., 2019; Sjodt et al., 2020; Taguchi et al., 2019; Zapun et al., 2012). aPBPs work independently from these two machineries, presumably filling in gaps and/or densifying the PG mesh (Cho et al., 2016; Straume et al., 2021; Vigouroux et al., 2020). PG remodeling on the other hand involves hydrolytic activities that cleave specific bonds within the nascent PG, allowing for example daughter cell separation or insertion of new material into the PG network (Vollmer et al., 2008b).

In this work, we investigated PG synthesis and remodeling in the opportunistic human pathogen *Streptococcus pneumonia*e. This bacterium belongs to the morphological group of ovococci (*e. g*. streptococci and enterococci), whose shape results from septal and peripheral PG syntheses, which respectively allow cell division and cell elongation (Briggs et al., 2021; Zapun et al., 2008). In *S. pneumoniae*, the divisome and elongasome contain the bPBP2x/FtsW and bPBP2b/RodA pairs, respectively (Briggs et al., 2021; Perez et al., 2019; Philippe et al., 2014; Straume et al., 2017; Zapun et al., 2012). None of the three pneumococcal aPBPs (aPBP1a, aPBP2a and aPBP1b) is essential but inactivation of both aPBP1a a and aPBP2a is lethal (Paik et al., 1999). Furthermore, aPBP1a modifies septal PG synthesized by the PBP2x/FtsW divisome pair (Straume et al., 2020). PG hydrolases related to cell division include the amidase/endopeptidase PcsB, the glucosaminidase LytB, and the muramidase Pmp23/MpgB (Bai et al., 2014; Barendt et al., 2009; Bartual et al., 2014; De Las Rivas et al., 2002; Jacq et al., 2018; Taguchi et al., 2021). On the other hand, the muramidase MltG/MpgA works in conjunction with the elongasome (Tsui et al., 2016).

Because PG synthetic and hydrolytic activities happen in regions with nanometer-scale dimensions, investigating their dynamics requires super-resolution microscopy methods that break the light diffraction barrier (Schermelleh et al., 2019). Recently, the combination of click-chemistry bio-orthogonal labeling with dSTORM (direct Stochastic Optical Reconstruction Microscopy) and *in silico* modeling improved our understanding of cell morphogenesis in the ovoid-shaped bacterium *S. pneumoniae* (Trouve et al., 2021b, 2021a). Proteins in charge of septal and peripheral PG synthesis are recruited to mid-cell by the cytoskeletal protein FtsZ at the beginning of the cell cycle, and their activities are first confounded within an annular region whose nanometric dimensions approximate that of the ring formed by FtsZ (Jacq et al., 2015; Trouve et al., 2021b). Later on, two synthesis regions progressively separate, the inner one corresponding to septal PG synthesis and the outer one to peripheral PG (Fig. S1A) (Perez et al., 2021; Trouve et al., 2021b). Importantly for this work, the study of PG dynamics by dSTORM brought support to a mechanism in which PG synthesis by the divisome at the membrane invagination front forms a septal matrix that is split at its periphery by hydrolases to form the new cell hemispheres. Peripheral PG is inserted by the elongasome at the periphery of the septum, into freshly cleaved septal material or concomitantly with the cleavage (Fig. S1A) (Trouve et al., 2021b). In this model, septum remodeling through cleavage and incorporation of peripheral PG would thus be a key event allowing both cell separation and cell elongation. Because they modify septal PG, aPBPs probably also influence septum cleavage and/or peripheral PG insertion (Straume et al., 2020).

With regards to the mechanisms of septum remodeling, the membrane-associated coiled-coil protein DivIVA is thought to be a main player (Halbedel and Lewis, 2019; Hammond et al., 2019). In *S. pneumoniae*, DivIVA is found at the division site and the cell poles, and it interacts with many elongation and division proteins, including bPBP2x, bPBP2b and aPBP1a (Fadda et al., 2007; Fleurie et al., 2014; Hammond et al., 2019; Straume et al., 2017). Deletion of the *divIVA* gene results in chains of short pneumococcal cells, indicative of impaired cell separation and elongation (Fadda et al., 2003; Fleurie et al., 2014; Straume et al., 2017). In *Streptococcus suis*, DivIVA interacts with MltG/MpgA and its absence causes a cell elongation defect (Jiang et al., 2023). Altogether, these observations point to a functional link between DivIVA and cell growth, but its exact function remains unknown.

Here, we used state-of-the-art fluorescence microscopy methods to investigate the role of DivIVA in *S. pneumoniae* cell morphogenesis. Using conventional fluorescence microscopy and STED (stimulated-emission-depletion), we describe the localization of DivIVA on both sides of the division site. Importantly, by combining bio-orthogonal PG labeling, dSTORM imaging and *in silico* modeling to analyze PG synthesis in *divIVA* mutants, we reveal that the dynamics of septum remodeling displays two phases along the pneumococcal cell cycle. During these two phases, DivIVA promotes both splitting of the septum and peripheral PG synthesis, ensuring that these two activities are synchronous around the septal periphery and maintained all along the division cycle.

## Results

### DivIVA localizes as a double ring at constricting division sites

Several studies have shown that DivIVA localizes at the division site in various bacterial species and in some cases at the cell poles, including in *S. pneumoniae* (Cramer et al., 2024; Fadda et al., 2007; Fleurie et al., 2014; Sutton et al., 2023). To further document the localization and dynamics of the pneumococcal DivIVA, we constructed a functional DivIVA-HT fusion at the endogenous *divIVA* locus (Fig. S2). Observation of DivIVA-HT using conventional fluorescence microscopy confirmed its septal and polar localization (Fig. 1A). Interestingly, it also showed an enrichment of the protein at mid- and late-division sites (constricting sites) compared with early-division sites (non-constricted sites) and cell poles (Fig. 1B). The localization of DivIVA at mid-cell is thus correlated with division stages during which the septum is being remodeled (Trouve et al., 2021b). STED microscopy further revealed the localization of DivIVA as a double band at mid-cell (Fig. 1C-D). These double bands, which correspond to the 2D projection of rings, are separated by a median distance of 93 ± 13 nm (n = 14).

**Figure 1.**
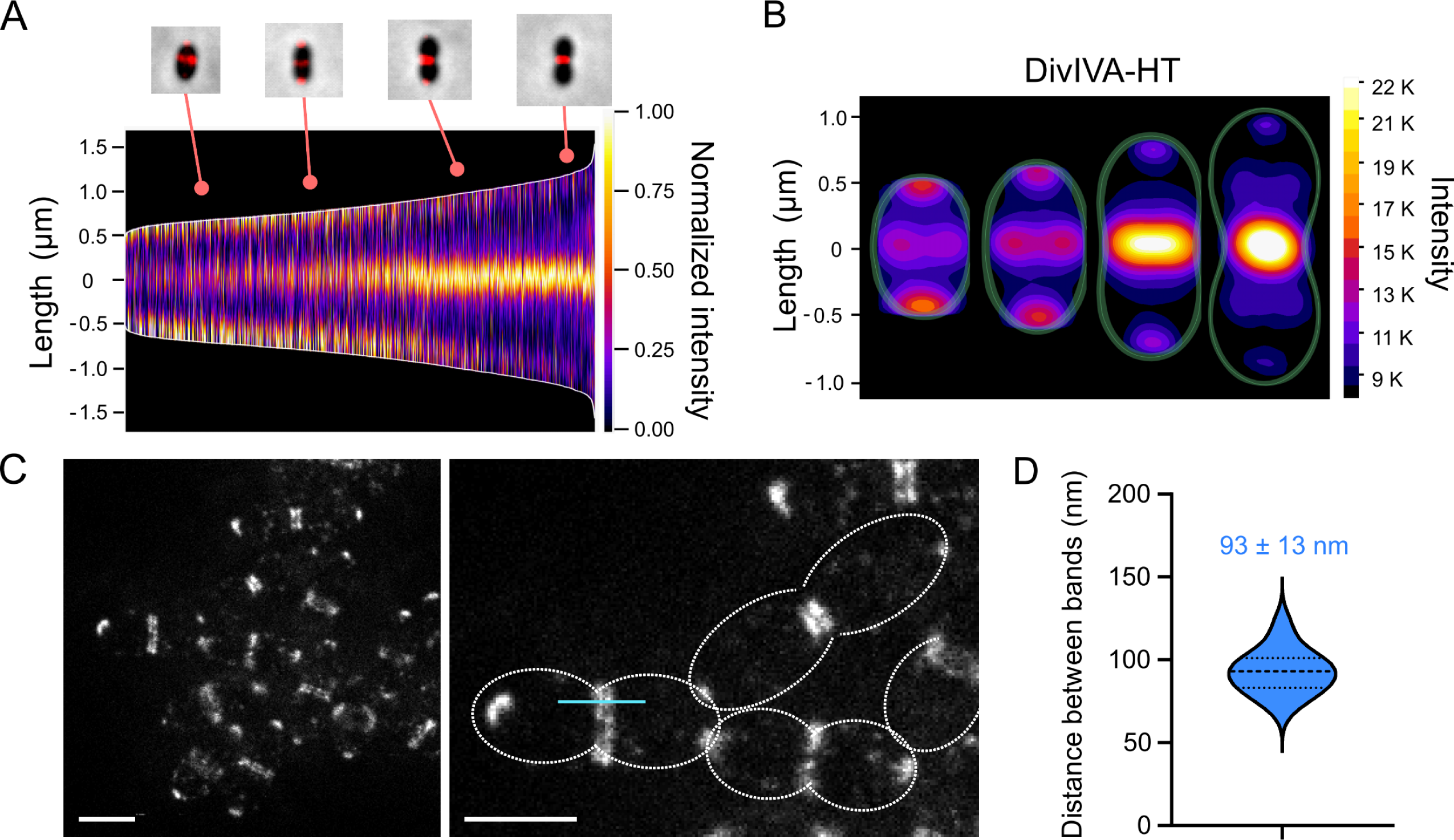
DivIVA localizes on both sides of constricting division sites in *S. pneumoniae*. Cells in exponential growth phase were incubated with HT-JFX646 ligand before fluorescence microcopy observation. **A.** Demograph showing the fluorescence intensity and diffraction-limited localization pattern of a DivIVA-HT fusion in 2,510 cells arranged according to their length. **B.** Heatmaps built from the localization patterns shown in (A) (n = 2,510), representing the average DivIVA-HT fluorescence intensity and localization along the cell cycle. Representative data are shown from two independent biological replicates. **C.** STED image showing the localization of DivIVA-HT as polar foci or as two bands flanking the division site. On the right panel, the cell contour is delineated with a dashed line and the blue line indicates a typical region used to generate intensity plot profiles. Scale bars, 1 µm. **D.** Violin plot showing the distribution of the distances separating DivIVA bands at midcell. The thin and thick dashed horizontal lines represent the quartiles (25^th^ = 83 nm; 75^th^ = 101 nm) and the median (93 nm) of two independent experiments, respectively. The median distance (n = 14) is indicated.

### DivIVA affects the dynamics of the pneumococcal cell cycle

In the absence of DivIVA, *S. pneumoniae* cells are still able to form complete septa but they do not cleave them efficiently, forming chains of more than 70 µm (Fig. 2A) (Fadda et al., 2007; Fleurie et al., 2014; Straume et al., 2017). To further characterize these defects, we analyzed the Δ*divIVA* strain by dSTORM following a short labeling pulse with the azido-D-Ala-D-Ala (aDA-DA) probe, which is incorporated into the growing PG (Trouve et al., 2021b, 2021a). The labeled cell chains display band-like patterns that result from the 2D projection of annular PG synthesis regions. The length of the labeled bands therefore measures the outer diameter of the pulse-labeled synthesis rings. Since they do not separate, the cells of the Δ*divIVA* strain could not be classified along the cell cycle as routinely performed with the wild-type (WT) strain (Fig. S1B) (Trouve et al., 2021b). Instead, we sought to classify the labeled PG synthesis sites. We observed an alternation of constricted and non-constricted labeling bands of various intensities (Fig. 2A). Interpreting the parental relationships between bands was complicated by the fact that a given site is not necessarily surrounded by sites of a single generation, but can also be bracketed by sites of different generations (Fig. 2B). We therefore based our classification on the diameter of the site considered with respect to that of its neighbors (Fig. 2C). We identified three classes of PG synthesis sites: the early-division class (E) harbors large diameters (> 800 nm) and no visible constriction, the mid-division group (M) exhibits visible constriction (*d_M_*) and neighboring sites that are even more constricted (*d* < *d_M_*), and the late-division group (L) includes constricted sites (*d_L_*) flanked by sites that are less constricted (*d* > *d_L_*). Our analysis revealed that most of the PG synthesis sites fall into the late-division group in the Δ*divIVA* strain, with 30%, 17% and 53% (n = 200) of the sites assigned to the early-, mid- and late-division classes, respectively (Fig. 2D). To compare these proportions with that of the WT strain, the cell categories reported previously were converted into categories of labeled bands by adding two early-division sites for each late division cell (Fig. S1B), since these were defined as displaying a constricted site surrounded by two new daughter sites (Trouve et al., 2021b). The division sites of the WT strain are thus distributed with proportions of early-, mid- and late-division of 68, 18 and 14%, respectively. Since cells were labeled during steady-state growth, the proportion of each labeled sites reflects the relative duration of the corresponding stage of the cell cycle. In the absence of DivIVA, the late-division stage is therefore longer than the earlier stages. Consistent with this observation and with a defect in septum splitting, the median diameters of the labeling sites in the Δ*divIVA* strain are higher than in the WT strain (Fig. 2E).

**Figure 2.**
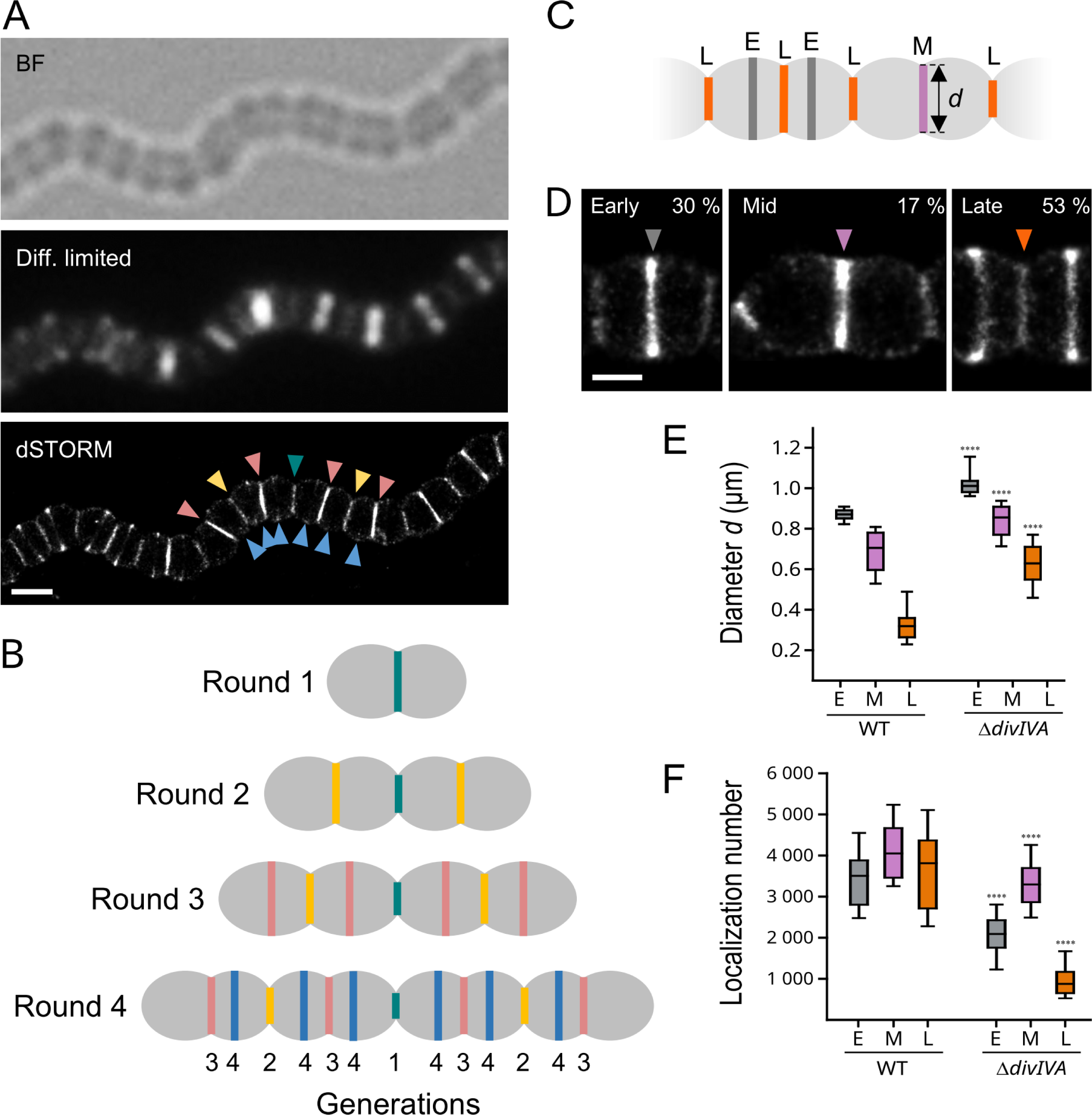
dSTORM analysis of PG synthesis in Δ*divIVA* cells. **A.** *S. pneumoniae* Δ*divIVA* cells in exponential growth phase were grown for 5 min in the presence of 2 mM aDA-DA, fixed and incubated with 35 µM DIBO-AF647 for fluorescent labeling. Bright field (BF), fluorescence diffraction-limited (Diff. limited) and reconstructed dSTORM images are shown. The arrowheads point to division sites of different generations, with a color code identical to that of panel B. Scale bar, 500 nm. **B.** Schematic illustration of the successive generations observed in Δ*divIVA* cell chains. **C.** Schematic diagram of pulse-labeled Δ*divIVA* cells lying along their longitudinal axis. The dark grey, purple and orange bands correspond respectively to early- (E), mid- (M) and late-division (L) pulse-labeling patterns, defined by their respective diameter (*d*). **D.** Zoom on representative dSTORM images of Δ*divIVA* cells displaying early-, mid- and late-division pulse-labeling patterns. Scale bar, 250 nm. **E.** Distribution of the diameters *d* among early- (*E*), mid- (*M*) and late-division (*L*) pulse-labeling patterns in wild-type (WT) (n = 116) and Δ*divIVA* (n = 200) cell populations. **F.** Distribution of the number of localizations among early- (*E*), mid- (*M*) and late- division (*L*) pulse-labeling patterns in wild-type (WT) (n = 116) and Δ*divIVA* (n = 200) cell populations. **E-F.** Data are represented with box plots showing the interquartile range (10^th^ and 90^th^ percentile), the median value and whiskers for minimum and maximum values. P-values from the U test of Mann-Whitney between WT and Δ*divIVA* data sets are indicated with quadruple asterisks when the observed difference is statistically significant (p-value < 0.0001).

Quantification of the number of localizations further shows that compared to the WT strain, less PG is synthesized in the absence of DivIVA (Fig. 2F). Whereas PG insertion remains robust along the whole cell cycle of WT cells, it dwindles to 27% of its maximum during late-division stages when *divIVA* is deleted. Also of note, the shortest labeled diameters measured in WT cells are at 150 nm, whereas the smallest diameter in the absence of DivIVA is about 280 nm (Fig. 2E). It is unclear whether cells fail to complete their cell division or if it continues at an extremely slow rate at which labeling is not detected.

### The pneumococcal cell cycle consists of two elongation phases

When scanning Δ*divIVA* cell chains, symmetrical patterns, sometimes of up to seven bands, can be found (Fig. 3). This number indicates that following a round of division, the cell cycles of the descendants can remain synchronized for two more rounds of division (see Fig. 2B). Labeling bands arising from a fourth generation are much less frequent, implying that the rate of cell cycle progression between cells vary sufficiently to nullify synchronization beyond three or four generations. We distinguished four typical symmetrical patterns. In one pattern (Fig. 3A), four long weakly-labeled bands (grey arrowheads, early stages) surround a short middle weakly-labeled band (central orange arrowhead, late stage) and two strongly-labeled bands of intermediate length (lateral orange arrowheads, late stage). In the second pattern (Fig. 3B), the central weakly-labeled band (central orange arrowhead, late stage) is flanked by four strongly-labeled bands (grey arrowheads, early stages), which sandwich two bands of smaller diameter (lateral orange arrowheads, late stages). The third pattern (Fig. 3C) is similar to the previous one, but the two small diameter bands show a weak labeling. A fourth pattern shows a series of weakly labeled bands (Fig. 3D).

**Figure 3.**
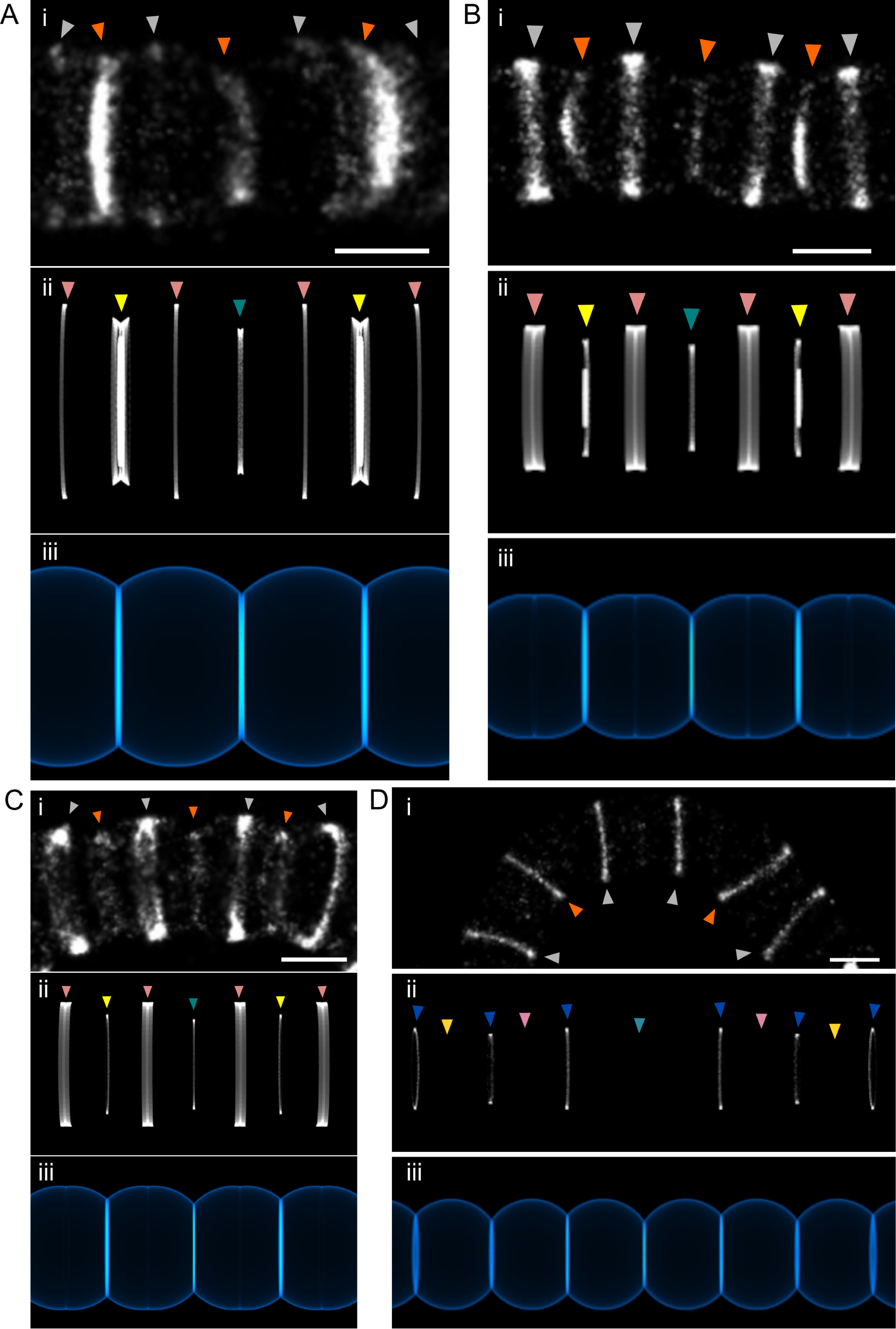
Symmetrical banding patterns are reproduced by a dynamic geometrical model of cell wall growth. *S. pneumoniae* Δ*divIVA* cells in exponential growth phase were grown for 15 min (**A** and **B**) or 5 min (**C** and **D**) in the presence of 2 mM aDA-DA, fixed and incubated with 35 µM DIBO-AF647 for fluorescent labeling. Reconstructed dSTORM (i), simulated labeling patterns (ii) and corresponding simulated cell wall (iii) images are shown. Scale bars, 500 nm. The same model parameters given in Table 1 were used for all the patterns, while the timing and duration of the labeling pulse were changed to generate each pattern (see Fig. S3).

**Table 1.** Parameters of the geometrical model used simulate the cell wall growth.

| strain | | WT<sup>a</sup> | $\Delta divIVA$ |
| --- | --- | --- | --- |
| Diameter | $D$ ($\mu\text{m}$) | 0.85 | 0.98 |
| Length | $L$ ($\mu\text{m}$) | 1.05 | 0.91 |
| Elliptic ratio | $E = L/D$ | 1.24 | 0.93 |
| Start of next generation as a fraction of the generation time | $N$ | 0.9 | 0.68 |
| Acceleration factor | $A$ | 0.9 | 0 |
| Septum closure as a fraction of generation time | $C$ | 0.84 | 0.68 |
| Time of slowing elongation as a fraction of generation time | $T_s$ | n/a | 0.58 |
| “Brake” factor | $B$ | n/a | 0.13 |
| Fraction of complete length at which all processes stop | $S$ | n/a | 0.8 |

<sup>a</sup> Reported from (Trouve et al., 2021b).

To understand the dynamics of PG synthesis underlying the observed dSTORM patterns, we previously developed a dynamic geometrical model of cell wall expansion. This model was parametrized using measurements taken on experimental data and could successfully recapitulate the labeling patterns of the WT strain (Fig. S1B) (Trouve et al., 2021b). Interestingly, to reproduce the patterns obtained with the Δ*divIVA* strain, this model had to be refined. Whereas modeling data from WT strains required an acceleration of the elongation rate to account for the thickening of the labeling bands along the cell cycle (Trouve et al., 2021b), the statistics of the band diameters and number of localizations measured for the Δ*divIVA* mutant (Fig. 2E-F) suggested a deceleration of the cell elongation. Different functions were trialed to model a deceleration of the elongation rate, but the morphology of the Δ*divIVA* strain and the labeling patterns could best and most simply be mimicked with two successive phases of constant elongation rates. Two additional parameters were thus added to the model to define when the second phase starts, and how reduced is its rate compared to the first phase. The absence of small labeled rings was modeled by a complete stop of the cell cycle at a fraction of the full cell length. The other parameters of the model remain. Regarding the cells, they are the length/diameter ratio, the time of completion of the septum closure and the time of the start of the next generation. Regarding the labeling patterns, the parameters are the start and the length of the pulse. In contrast to the WT strain for which the values of the parameters were obtained by fitting of the model functions to experimental measurements (Trouve et al., 2021b), the number of measurable, well-defined symmetrical patterns of the different types was not sufficient to perform meaningful fitting. Instead, the values of the parameters were manually adjusted to best represent the Δ*divIVA* observations. The modeled labeling patterns and cell wall representations are shown alongside dSTORM images in Fig. 3. The chosen values are given in Table 1. Graphs of the modeled evolution of the cell length, outer and inner diameter of the septum with the chosen parameters are given in supplementary Fig. S3. The first pattern (Fig. 3A) is produced when the labeling takes place when the second generation (yellow arrowheads) is producing most of its septum whereas the septum of the first generation (teal arrowhead) has already closed and the third-generation division has just started (pink arrowheads) (see Fig. S3 for the position of the green labeling pulse). In the second pattern (Fig. 3B), the pulse starts slightly later, while septum synthesis of the second generation is already well advanced (Fig. S3, orange labeling pulse). To generate the third pattern (Fig. 3C), a short pulse occurs as the third division cycle is in the first (fast) phase, whereas the second- and first-generation cycles are in the second slower phase and after septal closure (Fig. S3, cyan labeling pulse). The weak labeling is the result of slow peripheral PG insertion. The fourth pattern (Fig. 3D) appears when the labeling pulse occurs while three successive divisions have closed their septum and ceased to incorporate PG material but a fourth division (blue arrowheads) has just started (Fig. S3, yellow labeling pulse). The experimental data acquired on the Δ*divIVA* strain are thus consistent with a biphasic model of the pneumococcal cell cycle in which the elongation rate is reduced during the second phase.

### Septal synthesis is maintained but septum splitting and peripheral synthesis are impaired in the absence of DivIVA

To further analyze the dynamics of PG synthesis in the Δ*divIVA* strain, we analyzed tilted cells in which the radial architecture of the labeled regions can be observed (Fig. 4A). To account for the difference in cell diameter between the WT and Δ*divIVA* strains, we normalized the width of the septal plate (measured as the distance *r*, Fig. 4B) to the mean value of early-stage labeling diameters (*d(E)_WT_* = 850 nm; *d(E)_ΔdivIVA_* = 980 nm), as those have not yet initiated constriction. At mid- and late-division stages, the *r/d(E)* ratio is greater than in the WT strain (Fig. 4C), consistent with active septal PG synthesis and defective septum splitting in Δ*divIVA* cells. In this orientation, WT cells at mid- and late-division stages show inner and outer labeling rings, respectively corresponding to septal and peripheral PG synthesis (Fig. 4A, patterns *d* to *f*) (Trouve et al., 2021b). Cells harboring these concentric patterns represent 4% (n = 148) of the whole WT cell population. In Δ*divIVA* cells, similar concentric patterns are observed in a larger proportion of the cell population (15%, n = 148) (Fig. 4A, patterns *c* to *f*), which is consistent with impaired septum splitting. Indeed, since septal PG is synthesized at the leading edge of the growing septum and peripheral PG is synthesized at its periphery, deficient cleavage of the septum favors the separation of the two sites of synthesis. The experimental labeling patterns were correctly reproduced when septal synthesis was maintained until septum closure, and the rate of septum splitting and concomitant peripheral PG insertion was severely reduced in a second phase of the cell cycle (Fig. 4A).

**Figure 4.**
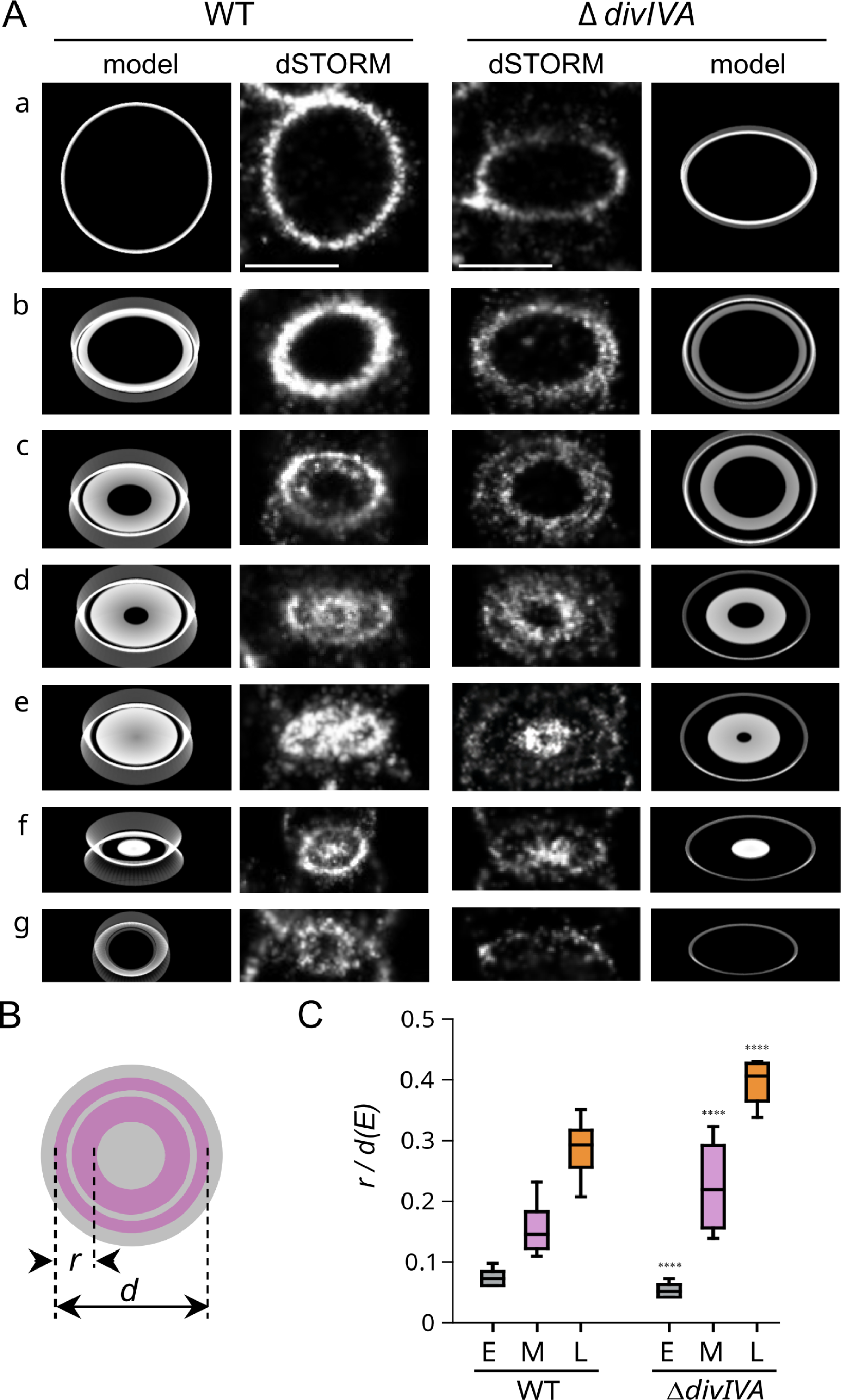
PG synthesis observed in tilted cells. **A**. Typical dSTORM localization patterns of pulse-labeled PG in cells oriented vertically or at an angle in an agarose pad. Early- to late-division stages are shown from top to bottom panels (*a-g*). Exponentially growing wild-type (WT) and Δ*divIVA* cells were incubated for 5 min with aDA-DA, fixed, and labeled with DIBO-AF647 before being trapped below an agarose pad. Scale bars, 500 nm. **B.** Schematic diagram representing a mid-division cell oriented perpendicular to its longitudinal axis. This orientation allows observing the septal (inner ring) and peripheral (outer ring) PG synthesis sites and measuring the radial width (*r*) of the labeled region. **C.** Distribution of the radial width of labeling *r*, normalized to the median largest diameter *d(E*) in early- (E), mid- (M) and late-division (L) pulse-labeling patterns in WT (n = 170) and Δ*divIVA* (n = 149) cell populations. Data are represented with box plots showing the interquartile range (10^th^ and 90^th^ percentile), the median value and whiskers for minimum and maximum values. The quadruple asterisks represent a statistically significant difference between WT and Δ*divIVA* datasets (U test of Mann-Whitney, p-value < 0.0001).

The dynamics of septum splitting and peripheral synthesis, or in other words cell elongation, were previously documented for WT strains using pulse-chase labeling experiments (Trouve et al., 2021b). Pulse-labeled PG is remodeled during the chase period, resulting in pulse-chase localization patterns that are wider than pulse patterns. In our model of pneumococcal PG synthesis, the septal PG is rapidly cleaved during early division, so that it is repositioned as hemispheric wall almost immediately after synthesis (Fig. S1A) (Trouve et al., 2021b). In a WT strain, when this stage (E) is observed in a pulse-chase experiment, the pulse-labeled PG splits into two bands during the chase, due to synthesis of unlabeled PG at mid-cell (Fig. 5A, grey arrowhead). At mid- (M) and late (L) division stages, septum cleavage lags behind septal PG synthesis, resulting in septum formation. In a pulse-chase experiment, a labeled septum is formed during the pulse and is cleaved during the chase. The cell wall that is repositioned laterally during the chase thus contains a mixture of labeled septally-synthesized PG and unlabeled peripheral PG (Trouve et al., 2021b). Such remodeling appears as “butterfly-like” patterns (Fig. 5A, pink and orange arrowheads). Throughout the cell cycle, the rate of septal cleavage and peripheral PG insertion can be assessed by measuring the “width of the chase” (*W_CHASE_*), which is the distance separating the external borders of the labeled patterns in pulse-chased cells (Fig. 5B).

**Figure 5.**
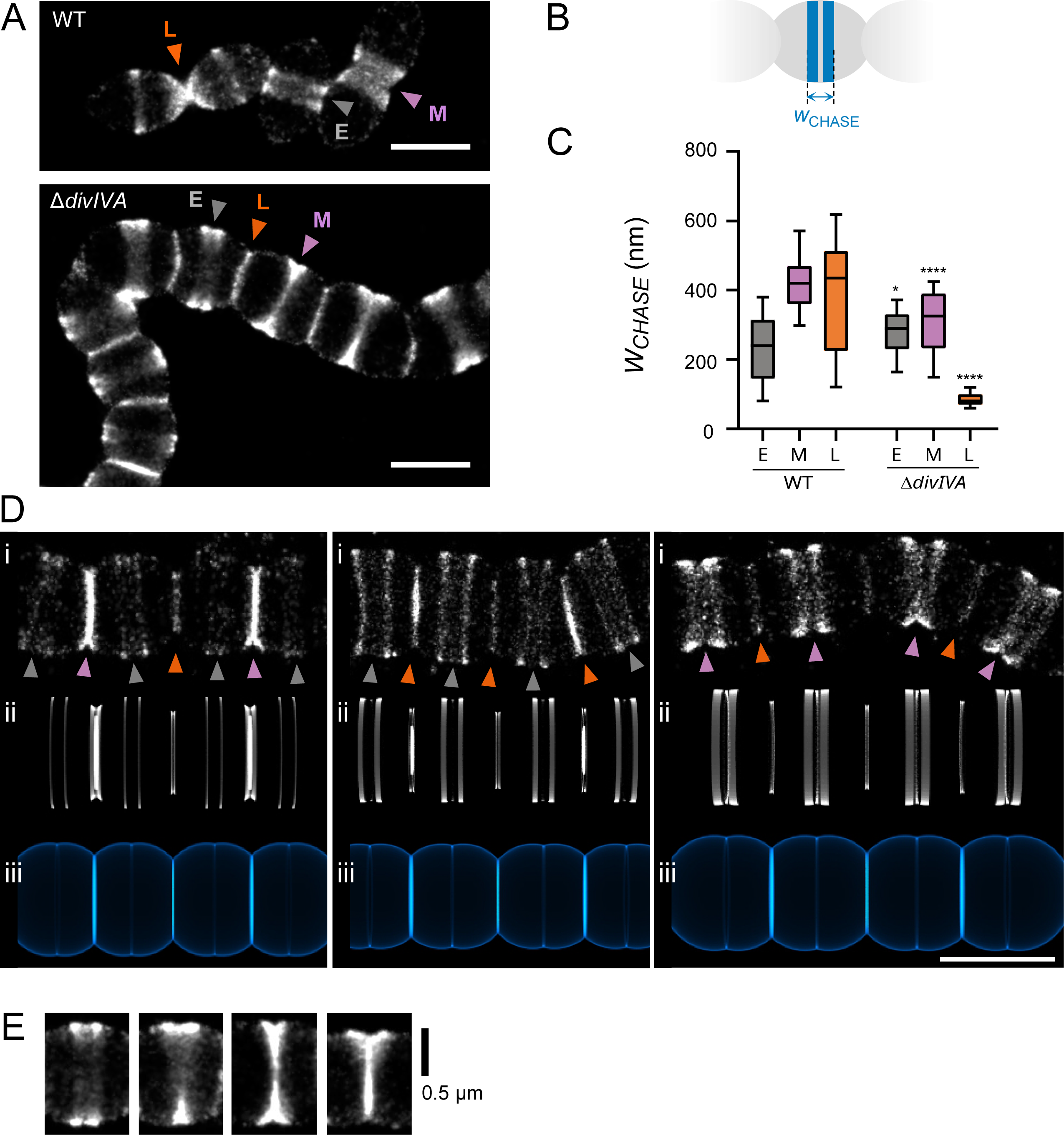
Pulse-chase labeling patterns of Δ*divIVA* cells. **A.** Representative dSTORM images of pulse-chase localization patterns in wild-type and Δ*divIVA* cells. Exponentially growing cells were incubated for 5 min with aDA-DA (pulse), grown for 15 min in the absence of the probe (chase), fixed, and labeled with DIBO-AF647 before dSTORM data collection. The grey, violet and orange arrowheads point to early-, mid- and late-division sites, respectively. Scale bars, 1 µm. **B.** Schematic representation of a pulse-chase labeled cell harboring two bands at an early-division site. The distance (*W_CHASE_*) separating the longitudinal outer limits of the labeling pattern is shown. **C.** Distributions of the *W_CHASE_* of the labeling patterns in early- (E), mid- (M) and late-division (L) sites (WT, n(*E*) = 89, n(M) = 68 and n(L) = 12; Δ*divIVA*, n(E) = 43, n(M) = 34 cells and n(L) = 101). Data are represented with box plots showing the interquartile range (10^th^ and 90^th^ percentile), the median value and whiskers for minimum and maximum values. P-values from the U test of Mann-Whitney between different data sets are indicated with single or quadruple asterisks when the observed differences are statistically significant with p-values < 0.01 or < 0.0001, respectively. **D.** Typical symmetrical dSTORM localization patterns of pulse-chase labeled PG in Δ*divIVA* cells. Earlier to later division stages are shown from left to right panels. Panels (i), (ii) and (iii) are dSTORM images, simulations of the labeling patterns and corresponding cell wall, respectively. Scale bar, 1 µm. **E.** dSTORM images of asymmetrical pulse-chase labeling patterns found with ΔdivIVA cells.

During early division, the median width of the chased region in Δ*divIVA* cells (*W_CHASE_(E)* = 270 nm, n = 119) is rather similar to that of the WT strain (*W_CHASE_(E)* = 260 nm, n = 123) (Fig. 5C). By contrast, *W_CHASE_*is reduced in the absence of DivIVA at mid- (*W_CHASE_(M)* = 280 nm, n = 57) and in particular at late-division (*W_CHASE_(L)*= 80 nm, n = 162) when compared to the WT strain (*W_CHASE_(M)* = 415 nm, n = 122; *W_CHASE_(L)*= 430 nm, n = 119). These measurements confirm that elongation slows down dramatically during the cell cycle in the absence of DivIVA, whereas it accelerates in the WT strain. The geometrical model correctly reproduces the observed chased patterns (Fig. 5D). Only sites that were labeled early during the first phase of rapid PG synthesis show significant parting of split labeled bands. Sites that were labeled late during the first (rapid) phase or during the second (slow) phase show little splitting of the bands.

In addition to the double-band patterns described above, 10% of the Δ*divIVA* early-division sites (n = 301) showed asymmetric separation of the labeled region, resulting in “Y-like” localization patterns (Fig. 5E). This proportion is probably underestimated since it only includes “Y-like” patterns correctly oriented in the XY plane and does not consider similar patterns oriented in the Z axis. Asymmetric pulse-chase labeling patterns are never observed in early-division WT cells, in which pulse-labeled regions are always chased as parallel bands (Fig. 5A). In the absence of DivIVA, the presence of Y-like labelings indicate that septal splitting and associated peripheral PG synthesis are non-uniformly distributed around the division site.

### The contribution of the elongasome and aPBPs to cell elongation is modified in the absence of DivIVA

*S. pneumoniae* cell elongation is known to rely on peripheral PG synthesis by the elongasome, which is formed by the RodA/bPBP2b pair (Perez et al., 2021; Straume et al., 2017; Vollmer et al., 2019). On the other hand, given that peripheral PG is inserted into cleaved septal regions and that septal PG is modified by aPBPs (Straume et al., 2020; Trouve et al., 2021b), it is possible that aPBPs influence peripheral PG insertion and thus cell elongation. Since the later process is impaired in Δ*divIVA* cells, we sought to investigate the interplay between DivIVA and the elongasome or aPBPs using pulse-chase experiments (Fig. 6A and 6C). Consistent with a previous study by Tsui et al, deletion of *pbp2b* in a WT background required the combined inactivation of *mltG*, which was obtained by mutating the catalytic tyrosine (Y488D mutation) (Tsui et al., 2016). Measurement of *W_CHASE_* was used to estimate the cell elongation. Compared to WT cells, *W_CHASE_* is reduced by a factor of 1.4, 1.5 and 1.6 at early-, mid- and late-division stages in Δ*pbp2b mltG_Y488D_* cells (Fig. 6B). In a WT background, bPBP2b-based peripheral PG synthesis is therefore a major contributor to all elongation phases. Interestingly, cell elongation is also diminished in mid- to late-division stages in the absence of aPBP1a or aPBP2a.

**Figure 6.**
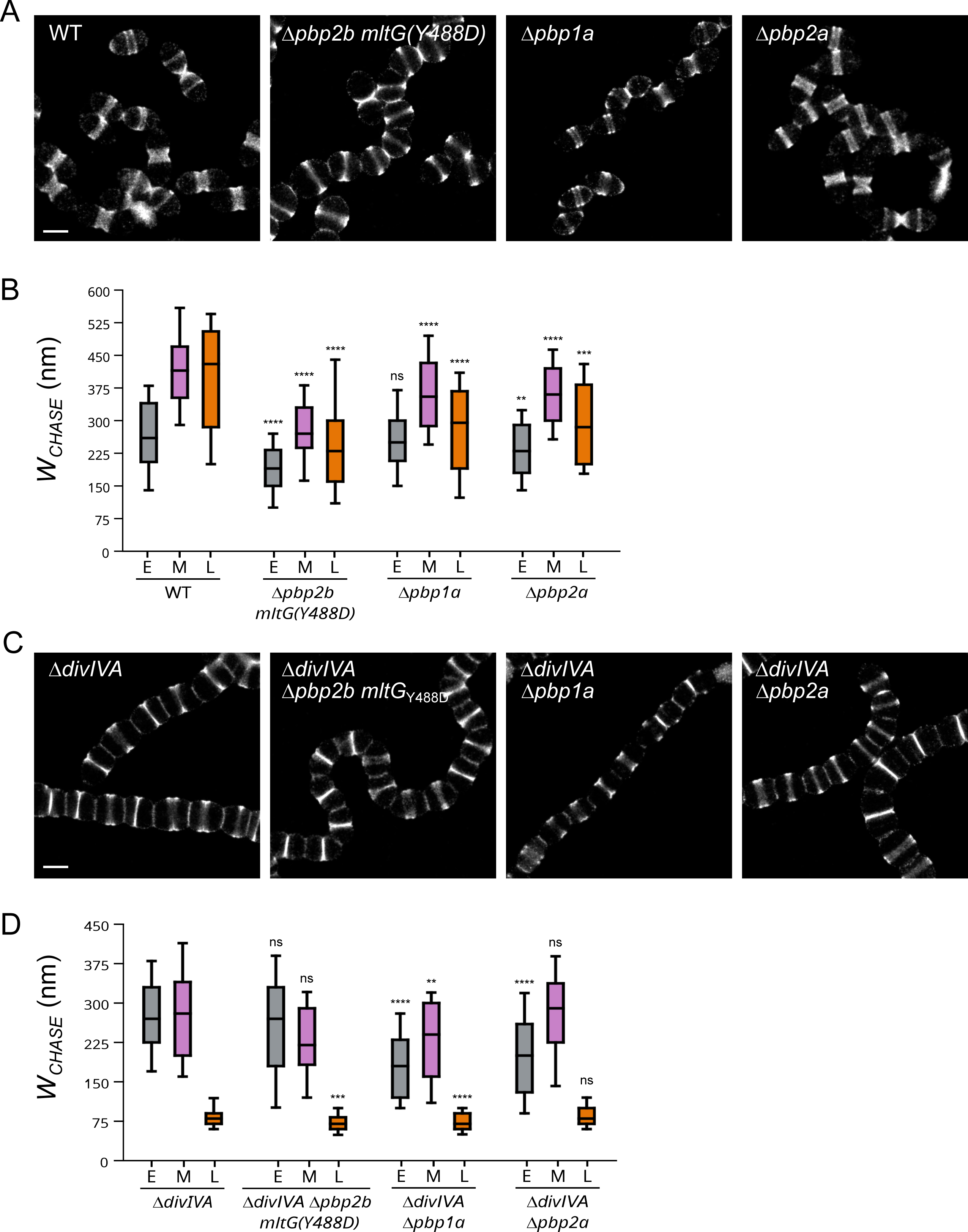
Effect of *pbp* gene deletion on Δ*divIVA* pulse-chase labeling patterns. **A.** Fields of wild-type (WT), Δ*pbp2b mltG(Y488D)*, Δ*pbp1a* and Δ*pbp2a* cells treated with the pulse-chase labeling protocol and observed by dSTORM. Scale bar, 1 µm. **B.** Distributions of *W_CHASE_* of the pulse-chase labeling patterns observed in panel A (WT, n(E) = 132, n(M) = 123 and n(L) = 46; Δ*pbp2b mltG*(Y488D), n(E) = 148, n(M) = 84 and n(L) = 65 cells; Δ*pbp1a*, n(E) = 140, n(M) = 116 and n(L) = 42 cells; Δ*pbp2a*, n(E) = 109, n(M) = 108 and n(L) = 40 cells). **C.** Fields of Δ*divIVA* mutant strains cells treated with the pulse-chase labeling protocol and observed by dSTORM. Scale bar, 1 µm. **D.** Distributions of *W_CHASE_* of the pulse-chase labeling patterns observed in panel C (Δ*divIVA*, n(E) = 120, n(M) = 57 and n(L) = 172 cells; Δ*divIVA* Δ*pbp2b mltG(Y488D)*, n(E) = 132, n(M) = 50 and n(L) = 120 cells; Δ*divIVA* Δ*pbp1a*, n(E) = 121, n(M) = 71 and n(L) = 109 cells; Δ*divIVA* Δ*pbp2a*, n(E) = 112, n(M) = 42 and n(L) = 146 cells). **B** and **D**. Data are represented with box plots showing the interquartile range (10^th^ and 90^th^ percentile), the median value and whiskers for minimum and maximum values. P-values from the U test of Mann-Whitney between different data sets are indicated with double asterisks when p-values < 0.01, triple asterisks when p-value < 0.001, quadruple asterisks when p-values < 0.0001, or with “ns” when no significant difference is observed (p-value > 0.05).

As reported previously (Fleurie et al., 2014), we could not delete the *pbp2b* gene in the Δ*divIVA* background, indicating that bPBP2b is still essential in the absence of DivIVA. As in a WT strain, we had to inactivate MltG to suppress the requirement for bPBP2b. However, in the Δ*divIVA* background, bPBP2b inactivation had a milder effect on cell elongation. Indeed, compared to Δ*divIVA* cells, *W_CHASE_* is either not reduced during early- and mid-division, or reduced by a factor of 1.2 at the late-division stage in Δ*divIVA* Δ*pbp2b mltG_Y488D_*cells (Fig. 6D). In the absence of DivIVA, the elongasome contribution to the cell elongation is thus diminished all along the cell cycle.

The comparison of the width of the chase between Δ*divIVA* and Δ*divIVA* Δ*pbp1a* cells shows a reduction in cell elongation at all stages of the cell cycle, especially in early- and late-division stages, upon *pbp1a* deletion (Fig. 6D). This is slightly different from the WT strain, in which the absence of aPBP1a has no effect on *W_CHASE_* during early-division (Fig. 6B). In a WT genetic background, the contribution of aPBP1a to early-elongation is probably masked by the strong contribution of the elongasome. The contribution of aPBP1a to early-division is therefore revealed by the absence of DivIVA, as the role of bPBP2b is minimal in this genetic background. In Δ*divIVA* cells, aPBP1a appears important for cell elongation all along the cell cycle.

The situation is somewhat different for aPBP2a. In the presence of DivIVA, aPBP2a mainly contributes to mid- and late-division stages (Fig. 6B). By contrast, in Δ*divIVA* Δ*pbp2a* cells, *W_CHASE_* does not decrease at these stages compared to Δ*divIVA* cells, indicating a decreased participation of aPBP2a to cell elongation in the absence of DivIVA (Fig. 6D). On the other hand, when compared to Δ*divIVA* cells, *W_CHASE_* is reduced by a factor of 1.4 during early-division in Δ*divIVA* Δ*pbp2a* cells. Here again, the role of aPBP2a during early cell elongation is therefore revealed in the absence of DivIVA.

In conclusion, the absence of DivIVA affects not only peripheral PG synthesis by the elongasome, but also the contribution of aPBP1a and aPBP2a to cell elongation.

## Discussion

DivIVA is anchored in the plasma membrane by an N-terminal motif composed of a hydrophobic residue flanked by positively charged amino acids (Choukate and Chaudhuri, 2020; Oliva et al., 2010). The central part of DivIVA forms coiled coils of varying length in different Gram-positive species. It assembles as a tetramer formed by the antiparallel association of two parallel coiled-coil dimers. Depending on the organism, there is an additional non-conserved C-terminal tail. Tetramers of DivIVA from *Bacillus subtilis* were found to assemble *in vitro* into large lattice-like structures (Oliva et al., 2010). Most importantly, DivIVA binds preferentially to negatively curved membrane regions, which accounts for its concentration at the cell poles and as two rings bordering the invaginating membrane, as previously reported for *B. subtilis* (Cramer et al., 2024), and established in this work for *S. pneumoniae*.

As a scaffold protein with an affinity for negatively curved membranes, DivIVA may have been used and reused by the mechanisms of evolution as a platform for processes that must occur at such places in bacteria. For examples, DivIVA directs apical growth in corynebacteria and mycobacteria (Kang et al., 2008; Letek et al., 2008), it participates indirectly to positioning of the division site in *B. subtilis* by interacting with the Min system (Eswaramoorthy et al., 2011), and it is involved in chromosome partitioning by positioning the ParB protein at poles (Giacomelli et al., 2022). In ovococci such as *S. pneumoniae*, the absence of DivIVA causes major morphological defects, as the Δ*divIVA* strain grows as chains of lentil-shaped cells, compressed in the direction of the long axis (this work and (Fleurie et al., 2014; Straume et al., 2017)). DivIVA is thus required for proper ovoid shape, but how does it fulfill its morphogenetic function?

Here, we have shown that in *S. pneumoniae*, DivIVA is more concentrated at the division site than at the cell poles, suggesting that it plays a major function at the sites of active PG synthesis. The current model proposes that the ovoid shape of *S. pneumoniae* results from a finely balanced and interconnected synthesis of septal and peripheral PG, respectively required for cell division and elongation. The latter is thought to result primarily from the action of the RodA/bPBP2b glycosyltransferase/transpeptidase couple associated with the hydrolase MltG/MpgA. Indeed, the absence of a functional elongasome (RodA/bPBP2b/MltG) engenders chains of shorter cells (this work and (Berg et al., 2013; Straume et al., 2017)), and our dSTORM data show that elongation is actually impaired at all stages of the cell cycle. Cells without DivIVA display a similar morphology, but we show in this case that elongation is only impaired at mid and late stages of the cell cycle. Surprisingly, in the same cells, the elongasome appears to be inactive in early and mid stages. Early elongation in Δ*divIVA* cells must therefore rely on another machinery than that of the elongasome. Observations made in aPBP mutants may shed some light on this. Indeed, in the absence of DivIVA, our data reveal a contribution of aPBP1a and aPBP2a to early elongation that is not detected in an otherwise WT background. Therefore, DivIVA seems to affect in some way the function of the elongasome and aPBPs, whose contribution to cell elongation seems to compensate each other during the early stage of the cell cycle.

Regardless of the enzymatic machineries involved, a major observation allowed by our dSTORM data is the existence of two cell growth phases during the pneumococcal cell cycle, as the absence of DivIVA does not affect early elongation (first phase) but dramatically impairs mid and late growth stages (second phase). More precisely, septation proceeds normally during the second phase but splitting of the septum and concomitant peripheral synthesis are drastically slowed down, leading to chains of shorter cells. From different attempts at simulating the labeling patterns with a geometrical model of the cell growth, it appears that the deceleration of septum splitting and peripheral synthesis must be abrupt rather that gradual. Later, it is possible that the elongation process ceases completely as no labeling of division sites of small diameter could be detected. Since long Δ*divIVA* cell chains still have extremities, the division of some rare cells may proceed to completion through septal splitting possibly decoupled from peripheral synthesis. Alternatively, long chains may be broken by the occasional lysis of some cells.

Also very interestingly, the resolution attained by dSTORM in pulse-chase labeling experiments has revealed Y-shaped labeling patterns in Δ*divIVA* cells, while pulse-labeled regions are always chased as parallel labeling rings in WT cells. These observations suggest that during the chase time, the elongation due to septal splitting and peripheral synthesis sometimes progresses at different speeds along the perimeter of the septum in the Δ*divIVA* strain, whereas the rate of elongation is identical around the septal circumference in WT cells.

Our detailed dSTORM localization of PG assembly in a *divIVA* deletion strain thus shows that DivIVA is necessary i) to maintain elongation passed a threshold time point during the cell cycle, and ii) to coordinate the related processes around the mid cell perimeter. How can DivIVA achieve such temporal and spatial regulation? Interestingly, the bi-phasic elongation process evidenced in this work is reminiscent of the bi-phasic dynamic behavior of DivIVA, which was shown to move in a directed manner around early-division sites and more diffusively around mid- and late-division sites (Perez et al., 2019). A hypothesis that would unify all the observations reported to date would thus be to consider that DivIVA is a physical sensor of membrane curvature, which would guide its localization in regions where septum splitting and peripheral synthesis need to be stimulated.

Let us first consider how should evolve the force that favors septum splitting (Fig. 7A). The surface tension resulting from the turgor pressure ΔP is γ = ΔPR/4 where R is the radius of curvature. At the junction of the septal and peripheral surfaces, for every segment *s* of the circumference of the septum, this surface tension generates a force, tangential to the peripheral surface away from the septum with a value 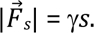 This force can be decomposed in a component parallel to the septal plane 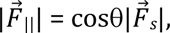 and a component perpendicular to the septal plane 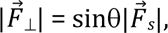 where θ is the angle between the new peripheral wall and the septal wall. This angle is π/2 at the start and increases towards π at the end of the division. Therefore, the component perpendicular to the septal plane 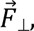 which should help tearing apart the septum, progressively vanishes as the division progresses. A good analogy of this phenomenon is the removal of an adhesive tape from a surface: it is easier if you pull perpendicularly to the surface than if you pull at a shallow angle. Thus, the turgor pressure and the cell geometry provide a force 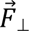 that can help septum splitting. The stretched septal PG at the periphery could be a better substrate for the cleaving enzyme.

**Figure 7.**
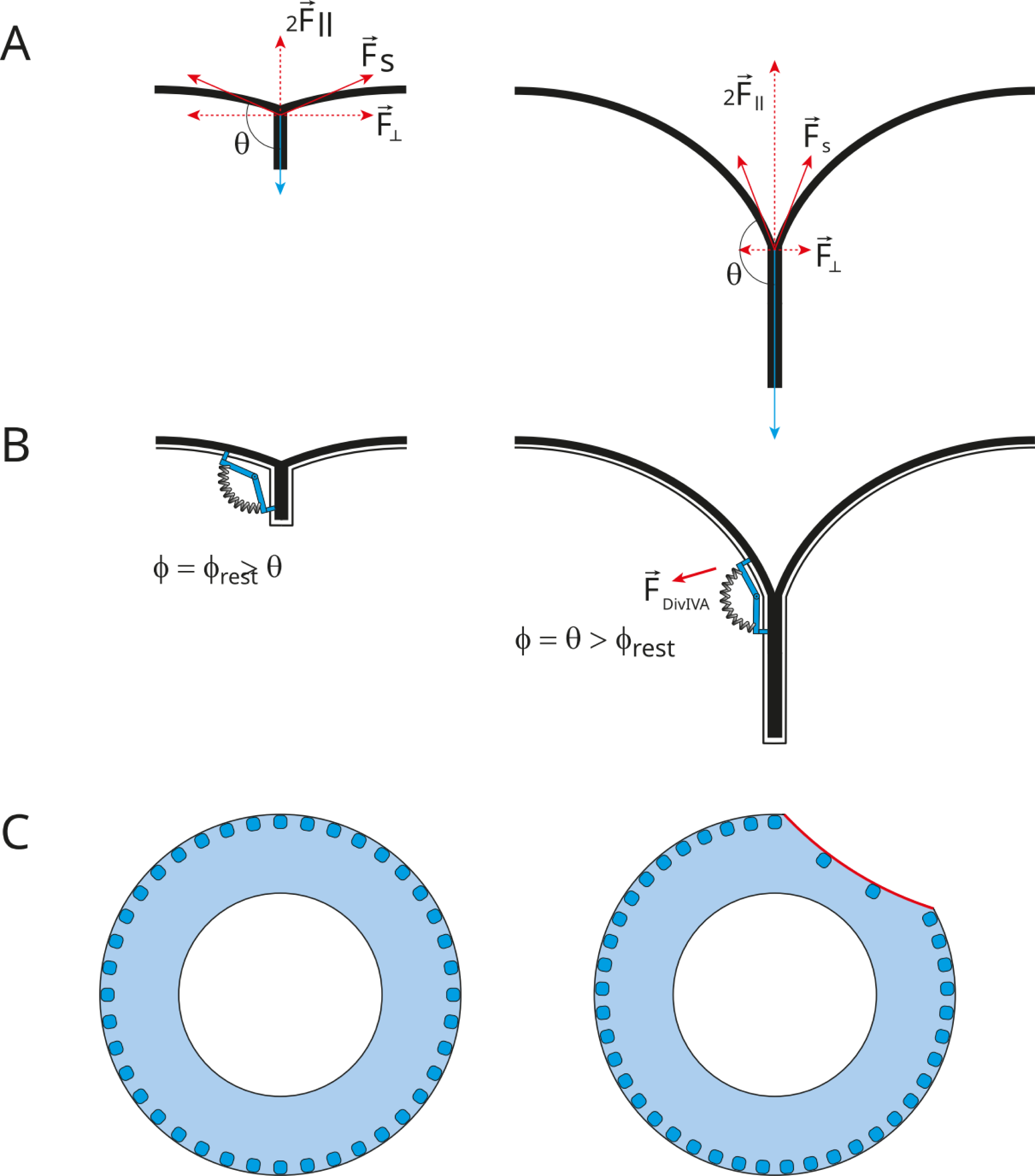
Model for the role of DivIVA in maintaining homogenous septum remodeling and peripheral PG synthesis in space and time. **A.** Schematic representation of the division site, with θ being the angle between the septal and peripheral walls and 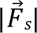 the tangential force exerted by the turgor pressure at the site of septum cleavage. Taking the septal plane as a reference, 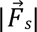 can be decomposed in a parallel 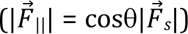 and a perpendicular 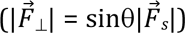 component. 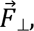 which contributes to septal PG cleavage, and decreases as the septum splits to generate new hemispheres. **B.** In virtue of its polymeric nature and affinity for negatively curved membranes, DivIVA might display a conformation of minimal energy defined by a resting angle (ϕ_rest_). Once ϕ_rest_ becomes smaller than θ, DivIVA stretching would generate a force that compensates the vanishing of the 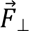 force, or activates the septum remodeling machinery. **C.** Schematic representation of how negative curvature affinity of DivIVA (blue dots) could ensure uniform progression of splitting and peripheral synthesis around the septum (blue ring). If splitting/peripheral synthesis occurs along one section of the septum (red), the membrane curvature along this section decreases, leading to a reduced concentration of DivIVA. The reduced amount of DivIVA would slow down the splitting/synthesis in this region in comparison to the rest of the periphery, thus restoring a uniform progression around the division site

The fact that in the absence of DivIVA, septum splitting and associated peripheral PG synthesis proceed normally during early division and then slows dramatically, suggests that a different or autonomous driving force exists at the beginning but is not present at the end of the process. The mechanical force 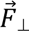 described above could fulfill this role since it diminishes necessarily during cell separation (Fig 7A). This waning force could reach a threshold where it is not sufficient and DivIVA would kick-in to stimulate further septum splitting. DivIVA, by virtue of its affinity for negatively curved membranes would be pre-localized exactly where required. In this scheme, DivIVA may be considered as an activator of septum splitting and the associated peripheral PG synthesis. How DivIVA achieves such activation remains mysterious. One common hypothesis is to propose that DivIVA acts as a platform protein that interacts directly or indirectly with proteins of the cell wall metabolism to concentrate them and/or activate them at specific locations. Based on the defective peripheral synthesis observed in Δ*divIVA* cells, elongasome proteins appear as putative partners of DivIVA. In agreement with this idea, DivIVA was shown to localize (but not activate) MltG/MpgA through a direct physical interaction in *Streptococcus suis* (Jiang et al., 2023). MltG/MpgA might thus use DivIVA as a localization platform. This hypothesis seems less likely for bPBP2b as it remains localized in the absence of DivIVA, but we cannot exclude that DivIVA induces local variations in bPBP2b concentrations that were not detected in the study by Fleurie et al (Fleurie et al., 2014). An alternative hypothesis would be that DivIVA might indirectly activate peripheral synthesis, again through a physical mechanism based on its negative-curve-affinity (Fig 7B). Let us assume two additional features for DivIVA: a resting conformational angle ϕ_rest_ in the longitudinal plane comprised somewhere between π/2 and π, and anchoring points to the cell wall at both extremities (through some relay proteins). At the beginning of the division, the DivIVA angle would be greater than that of the membrane at the periphery of the septum (ϕ = ϕ_rest_ > θ). Since θ increases as the septum is cleaved, at some point the DivIVA resting angle becomes smaller than that of the membrane (ϕ_rest_ < θ) and DivIVA would be stretched (ϕ = θ). In this new configuration, DivIVA experiences a tension that may be translated into an activation signal to interacting PG synthases/hydrolases, or it could transfer tension to the periphery of the septal PG itself, taking over the function of the diminishing splitting force 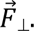 Note that the two hypothetical modes of action are not mutually exclusive.

If the affinity of DivIVA for the membrane increases with the curvature, and if septum splitting and peripheral synthesis are positively regulated by the presence of DivIVA, these two conditions would ensure an orderly symmetrical proceeding of the division around the cell circumference. If division were to occur faster in one region of the circumference (Fig. 7C right), the peripheral membrane curvature in that region would become smaller than in the rest of the cell circumference; the concentration of DivIVA would then decrease in this region of faster division and increase elsewhere, where it would stimulate the process. The sliding and modification of the local concentration of DivIVA along the septal circumference according membrane curvature would thus automatically adjust the amount of cell wall metabolism required to ensure the even progression of the division. In Δ*divIVA* cells, heterogeneous septum splitting and/or peripheral synthesis would not be corrected by DivIVA, resulting in circularly asymmetrical PG labeling profiles, observed as Y-shaped labeling patterns in our dSTORM images.

The biphasic dynamic behavior of DivIVA evidenced by Perez et al. can also be explained by its affinity for negatively curved membranes (Perez et al., 2019). When the septum is not yet formed, if FtsZ drives membrane invagination at the division site, places of maximal negative membrane curvature necessarily follow the localization of the Z-ring around the cell perimeter. The affinity of DivIVA for negative membrane curvature would result in DivIVA moving directionally around the mid-cell circumference, seemingly “following” FtsZ treadmilling during early division. Past the very early stage, a maximal negative curvature is established at the periphery of the septal plate, whereas FtsZ remains localized at the leading edge of the closing septum. Consequently, positioning of DivIVA would not depend on FtsZ treadmilling after the initial moments of the cell division process. At these later stages, the localization of DivIVA molecules around the septal periphery would depend on modulation of the negative curvatures resulting from septum splitting, as proposed above.

Although the mode of action of DivIVA remains to be elucidated, our work improves our understanding of the cell cycle in *S. pneumoniae* and highlights a regulatory function for DivIVA. It sets the ground for further investigation, which should include cellular studies of DivIVA dynamics, biophysical studies of DivIVA interaction with the membrane, as well as high-resolution structural studies in the cell, such as cryo-correlative light and electron microscopy (cryo-CLEM), to correlate the localization of DivIVA with the architecture of cellular ultrastructures.

## Materials and Methods

### Bacterial strains and growth conditions

The pneumococcal strains used in this study are listed in Table S1 and the strain construction is described in *Supplementary Methods*. For PG labeling, 50 µL of glycerol stocks, frozen at OD_600nm_ = 0.3, were used to inoculate 1 mL of BHI medium. Cultures were grown at 37°C in a static CO_2_ incubator, diluted twice into 10 mL of pre-warmed BHI medium (1/10^th^ volume) to reach steady-state growth, and treated for PG labeling when OD_600nm_ reached 0.3.

To localize DivIVA, cultures were inoculated at OD_600nm_ = 0.006 in C+Y medium and grown at 37°C to an OD_600nm_ of 0.3. These cultures were diluted into C+Y medium (1/50^th^ volume) and incubated at 37°C to an OD_600nm_ of 0.1. The cells were then incubated 10 min at 37°C with 20 nM HT-JFX646 ligand. For STED Microscopy, to fix the cells, 300 μL of culture were mixed with 100 μL of fixation solution (0.5 M KPO_4_ pH 7, 8% paraformaldehyde, 0.08% glutaraldehyde) and incubated for 15 min at room temperature followed by 15 min on ice. Cells were then washed twice with 300 μL of 0.1 M KPO_4_ and finally concentrated 3 times. 0.8 μL of the cell suspension was immobilized on a 1.2% C+Y agarose-coated microscopy slide with a cover glass (# 1.5) for cell imaging.

### Epifluorescence and STED microscopy

Epifluorescence snapshots were performed on a previously described setup, equipped with a 100 x objective and with laser excitation at 561 nm set at 30% of the maximum excitation power with exposure time of 500 ms (Billaudeau et al., 2017). Demographs showing the fluorescence intensity of DivIVA in relation to cell length was generated using MicrobeJ (version 5.13n), with specific parameters previously reported (Juillot et al., 2024): area (0.6-20 μm²), length (0.5-max μm), width (0.5-2 μm), circularity (0-1), curvature (0-max), sinuosity (0-max), angularity (0-0.25 rad), solidity (0-0.85 max), and intensity (0-max). Cells with regular morphology that were initially excluded from the segmentation due to proximity to other cells were included manually in the analysis. Adjacent ovoid cells were meticulously examined to ensure accurate classification as one late-division cell or two separated daughter cells. The resulting data were normalized as per MicrobeJ’s guidelines.

Heatmaps illustrating the average intensity of DivIVA-HT within cells was produced with MicrobeJ, using cell categorization parameters previously reported as follows (Juillot et al., 2024): from left to right, [SHAPE.length/SHAPE.width] ≤ 1.7; [SHAPE.length/SHAPE.width] > 1.7 and [SHAPE.length] ≤ 1.6; [SHAPE.length/SHAPE.width] > 1.7 and [SHAPE.length] > 1.6; [SHAPE.length/SHAPE.width] > 2 and [SHAPE.circularity] < 0.8; [SHAPE.length/SHAPE.width] > 2 and [SHAPE.circularity] < 0.7.

STED microscopy images were acquired with a STEDYCON (Abberior Instruments GmbH). Pulsed laser at 640 nm was used for fluorescence excitation. A pulsed 775 nm STED depletion laser was additionally used. The 640 nm excitation laser and the 775 nm depletion laser were set at 10% of their maximum power. The optical setup was connected to an AxioImager Z2 fluorescence microscope (Zeiss) with a 100× oil immersion PlanApochromat objective lens (NA = 1.46). Acquisition software was STEDYCON Smart control. Measurements were performed with a pixel dwell time of 10 μs.

### Preparation of PG labeling reagents

All starting materials were obtained from commercial sources and were used without further purification. NMR spectra were acquired on a Bruker Advance 400 MHz for ^1^H-NMR experiments and 100 MHz for ^13^C-NMR experiments. Chemical shifts are reported in ppm (*δ*) relative to the solvents: ^1^H *δ*(CD_3_OD) = 3.3 ppm, ^13^C *δ*(CD_3_OD) = 49.15 ppm. Accurate mass spectra were recorded on a time-of-flight (TOF) spectrometer (Waters, XEVO G2-S QTof) and on a LTQ Orbitrap XL spectromer with ElectroSpay Ionization (ESI, Thermo Scientific).

The aDA-DA probe (((*R*)-2-amino-3-azidopropanoyl)-D-alanine) was synthesized as previously described (Trouve et al., 2021b). Briefly, 2-(trimethylsilyl)ethyl (*tert*-butoxycarbonyl)-D-alaninate was obtained according to the Van Nieuwenhze’s process and the (*R*)-3-azido-2-((*tert*-butoxycarbonyl)amino)propanoic acid was obtained according to the Webb’s process (described for the L enantiomer) (Liechti et al., 2014; McAllister et al., 2011).

The N-Boc deprotection of 2-(trimethylsilyl)ethyl (*tert*-butoxycarbonyl)-D-alaninate was performed in a solution of 4 N HCl in dioxane for 3 h to give, after removal of solvent, trituration in dried ether, and drying under high vacuum (4 h), the corresponding salt in quantitative yield. This latter and the (*R*)-3-azido-2-((*tert*-butoxycarbonyl)amino)propanoic acid (1.4 equivalent) were dissolved in dry CH_2_Cl_2_. HATU (1.1 equivalent), di-isopropyl ethyl amine (DIPEA, 2.3 equivalent) was added and the mixture was stirred overnight. After removal of the solvent, ethyl acetate was added and the solution was washed with an aqueous solution of 10% citric acid followed by 5% NaHCO_3_. The combined organic layers were dried (MgSO_4_) and evaporated. After purification flash chromatography, the resulting 2-(trimethylsilyl)ethyl ((*R*)-3-azido-2-((*ter*-butoxycarbonyl)amino)propanoyl)-D-alaninate was obtained in 63% yield. The final deprotections were performed by addition of trifluoroacetic acid (TFA) at 0°C followed by stirring for 4 h at room temperature. The solvent was evaporated and dried under high vacuum (oil pump). The compound was purified by HPLC to yield ((*R*)-2-ammonio-3-azidopropanoyl)-D-alaninate 2,2,2-trifluoroacetate (aDADA) as a white solid.

Frozen aliquots of a 500-mM stock solution of aDA-DA in 100% DMSO were diluted to 10 mM in commercial DPBS (calcium- and magnesium-free, from ThermoFisher) and stored at −20°C for subsequent use.

Powder of DBCO-AF647 (Jena Bioscience) was resuspended at 10 mM into 100% DMSO, diluted to 500 µM into DPBS and stored at −20°C for subsequent use.

### PG labeling in *S. pneumoniae* cells

Cells in exponential growth phase (OD_600nm_ 0.3) were incubated for 5 min at 37°C in BHI medium containing 2 mM aDA-DA (PULSE labeling). For PULSE-CHASE labeling experiments, after the 5-min PULSE labeling with aDA-DA, cells were pelleted (7,000 × g, 5 min, 20°C) and resuspended into fresh, pre-warmed BHI medium. Incubation at 37°C was continued for 15 min. Following PULSE or PULSE-CHASE labeling, cells were fixed overnight on ice, in the culture medium supplemented with 0.5-X DPBS and 2% (v/v) paraformaldehyde (PFA). Following fixation, cells were pelleted, resuspended into DPBS containing 35 µM DBCO-AF647 and incubated for 1 h at 20°C (click reaction). Cells were then washed twice with 1 mL of DPBS and were resuspended into the dSTORM buffer, which contains 25 mM NaCl, 75 mM Tris-HCl pH 8.0, 10% (w/v) D-glucose, 100 mM cysteamine (β-mercaptoethylamine) and 1X GLOX mix [40 µg⋅mL^-1^ catalase from bovine liver (Sigma-Aldrich), 0.5 mg⋅mL^-1^ glucose oxidase type VII from *Aspergillus niger* (Sigma-Aldrich)]. Further details regarding the labeling procedure and alternative strategies can be found in (Trouve et al., 2021a).

### dSTORM data acquisition

To perform dSTORM on cells oriented along their longitudinal axis, the samples were mounted between a slide (High-precision, No. 1.5H, 24 × 50 mm, 170 ± 5 mm, Marienfeld) and a coverslip (High-precision, No. 1.5H, 22 × 22 mm, 170 ± 5 µm, Marienfeld) previously treated with ozone and eventually sealed with colorless nail polish.

To orient cells with a tilt angle, 8 µL of sample was loaded on a slide and covered with 18 µL of dSTORM buffer containing 0.9% (w/v) low-melting agarose. The slide was then turned over onto a coverslip. After solidification, the agarose pad was sealed with colorless nail polish.

dSTORM images were acquired at 27°C with a SAFe360 Abbelight PALM/STORM setup based on an Olympus IX83 inverted microscope equipped with a Oxxius laser combiner and two sCMOS Hamamatsu ORCA-Fusion cameras controlled via the NEO software. The emitted signal was collected by a 100x oil immersion apochromatic objective (UPLAPO100XOHR, Olympus) and then filtered by a band-pass filter FF01-698/70 (Semrock). First, diffraction-limited images were acquired by illuminating the sample at 640 nm with 1% of the maximal laser power (2 kW⋅cm^-2^). For single-molecule localization microscopy, the samples were exposed to a continuous 640 nm illumination with a constant laser power density of 2 kW⋅cm^-2^. dSTORM data were collected over 15,000 frames recorded with an exposure time of 50 ms. The blinking density was maintained all along the data collection by ramping up a 405-nm laser from 0% to 15% of a maximal value of 50 W⋅cm^-2^ For homogenous excitation of the sample, the field of excitation parameter in the NEO software was adjusted to cover the region of interest (ASTER illumination) (Mau et al., 2021).

### dSTORM image reconstruction

Data processing was performed with the ThunderSTORM plugin in ImageJ/FIJI (Ovesný et al., 2014; Schindelin et al., 2012; Schneider et al., 2012), including determination of the localization precision and quantification of the number of localizations. Potential drift was corrected with the cross-correlation tool in ThunderSTORM (Ovesný et al., 2014). Image rendering was achieved through normalized Gaussian blurring with a reconstructed pixel size of 10 nm. The spatial resolution of the reconstructed dSTORM images was calculated using the FIRE ImageJ/FIJI plugin (Nieuwenhuizen et al., 2013). Further details for dSTORM data acquisition and image reconstruction can be found in (Trouve et al., 2021a).

### Measurement of the labeling patterns

Measurements of the labeled regions were performed in ImageJ (Schneider et al., 2012). The values of the diameter *d* and of the width of the chase *W_CHASE_* (as defined in figures 2C and 5B) were obtained by plotting the signal intensity along a line positioned over the labeling of interest and measuring the distance between the points at half the maximal intensity.

For radial labeling patterns (in vertical or angled bacteria), the diameters *d_out_* and *d_in_* used to calculate the radial thickness *r* of the labeled region, as defined in figure 4B, were measured as above along a line positioned across the longest axis of the elliptic patterns.

### Geometrical model of cell growth and labeling

The basis of the geometrical model was similar to that developed previously for WT strains (Trouve et al., 2021b). The animated geometrical model was implemented using the free open source 3D animation software Art of Illusion© by Peter Eastman. The mature cell was modeled as an ellipsoid of length *L* and diameter *D*. Taking the longitudinal axis as the *x* axis, the equator as its origin, and *L* the length of a mature single cell, the radius of the cell around this axis (*R_out_*) is given by the elliptic curve:

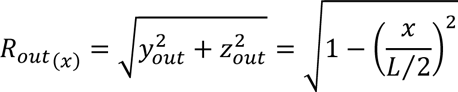

Taking *l* as the distance between two parting equators, as the new cell length *l/2* grows along *x* with time, *R_out_*at *x* = *l/2* is the radius of the septal outer edge circle (*R_out_*= *d_out_*/2). Taking the length *C,* at which the septum is closed, the radius of the septal inner edge circle (*R_in_* = *d_in_*/2) at *x* = *l*/2 is given by:

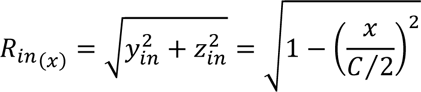

That points on the inner septal edge follow an elliptic path towards the point of closure is an assumption that was found to match the observations of the WT strains. We have set *R*_max_ = *D*/2 = 1, so that in the above equations, *L*/2 takes the numerical value of the elliptic factor *E = L/D*.

To account for the acceleration of the growth along the *x* axis observed in WT strains, *l/2* depends on time according to:

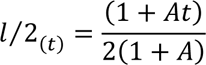

where *A* is an acceleration factor. Note that in this equation, *t* and *A* are dimensionless and *t* is expressed as a fraction of the generation time, which is defined as 1. The (1+*A*) denominator is a normalization factor that ensures that a division is completed at *t* = 1.

Since the acceleration was not observed in the Δ*divIVA* strain, *A* was set to 0 to describe the deletion phenotype. Instead, three additional parameters were introduced in the model. A time point after the initiation of the division at which the cell elongation slows down (*T_s_*), and the factor by which the elongation rate is diminished (“brake” factor *B*). When *B* is set to 1, growth continues unchanged after *T_s_* at the normalized rate of 1 and the division cycle is completed at *t* = 1. If *B* is set as a fraction of 1, the cell elongation continues at rate *B* after *T_s_*, and the division ends at *t* = *T_s_* + (1 − *T_s_*)/*B*, which is greater than 1.

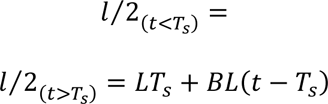

The third parameter that was added determines a point at which all processes stop. For practical reason of programming in Art of Illusion©, this event was defined as a fraction (*S*) of the length that the cell would have attained without growth arrest, rather than as a time.

The start and length of the labeling pulse can also be defined. In contrast to the WT strains, where the evolution of the various distances (length *l*, inner and outer septal diameter *d_in_* and *d_out_* could be fitted to measurements to extract values of the model parameters, the parameter describing the Δ*divIVA* strain were manually adjusted to best reproduce the observations, due to the greater heterogeneity of the cell sizes and morphologies. Nevertheless, all the simulated labeling patterns presented were generated with the same set of parameters, which are given in Table 1.

## Quantification and Statistical Analysis

Statistics were performed using Excel and figures were prepared using LabPlot2. The statistical details describing the quantification and measurements of localization patterns (including the definition of center, dispersion and precision of measures, the statistical tests used, the exact value of the *n* number of analyzed cells) can be found in the *Results* section and in the figure legends.

## Materials and Data Availability

There are restrictions to the availability of the aDA-DA probe. Due to the time required for chemical synthesis, this material will be available through a collaboration with Yung-Sing Wong.

The datasets generated during this study will be made available on the Zenodo platform upon acceptation of the manuscript [accession # To be named/zenodo.org].

## Acknowledgments

We thank members of the Morlot and Grangeasse laboratories for advice and encouragement, O. Glushonkov and J.P. Kleman for advices and support regarding dSTORM. Support for this work comes from the Agence Nationale de la Recherche (ANR-16-CE11-0016 and ANR-23-CE11-0029 to CM, ANR-19-CE15-0011 to CG, Labex ARCANE ANR-17-EURE-0003 to YSW, and ANR-17-CE13-0031 to RCL and NC). This work used the platforms of the Grenoble Instruct-ERIC centre (ISBG; UMS 3518 CNRS-CEA-UGA-EMBL) within the Grenoble Partnership for Structural Biology (PSB), supported by FRISBI (ANR-10-INBS-05-02) and GRAL, financed within the University Grenoble Alpes graduate school (Ecoles Universitaires de Recherche) CBH-EUR-GS (ANR-17-EURE-0003). IBS acknowledges integration into the Interdisciplinary Research Institute of Grenoble (IRIG, CEA).

## Author contributions

JT, AZ, RCL, NC, YSW, CG and CM designed research; JT, AZ, LB, DJ, AP, CF and MB performed experiments; JT, AZ, DJ, NC, YSW, CG and CM analyzed data; JT, AZ and CM wrote the manuscript; JT, AZ, RCL, NC, YSW, CG and CM revised the manuscript.

## Inclusion and Diversity Statement

While citing references scientifically relevant for this work, we also actively worked to promote gender balance in our reference list.

