## Supplementary Information for "DivIVA controls the dynamics of septum splitting and cell elongation in *Streptococcus pneumoniae*"

**Supplementary Figures**

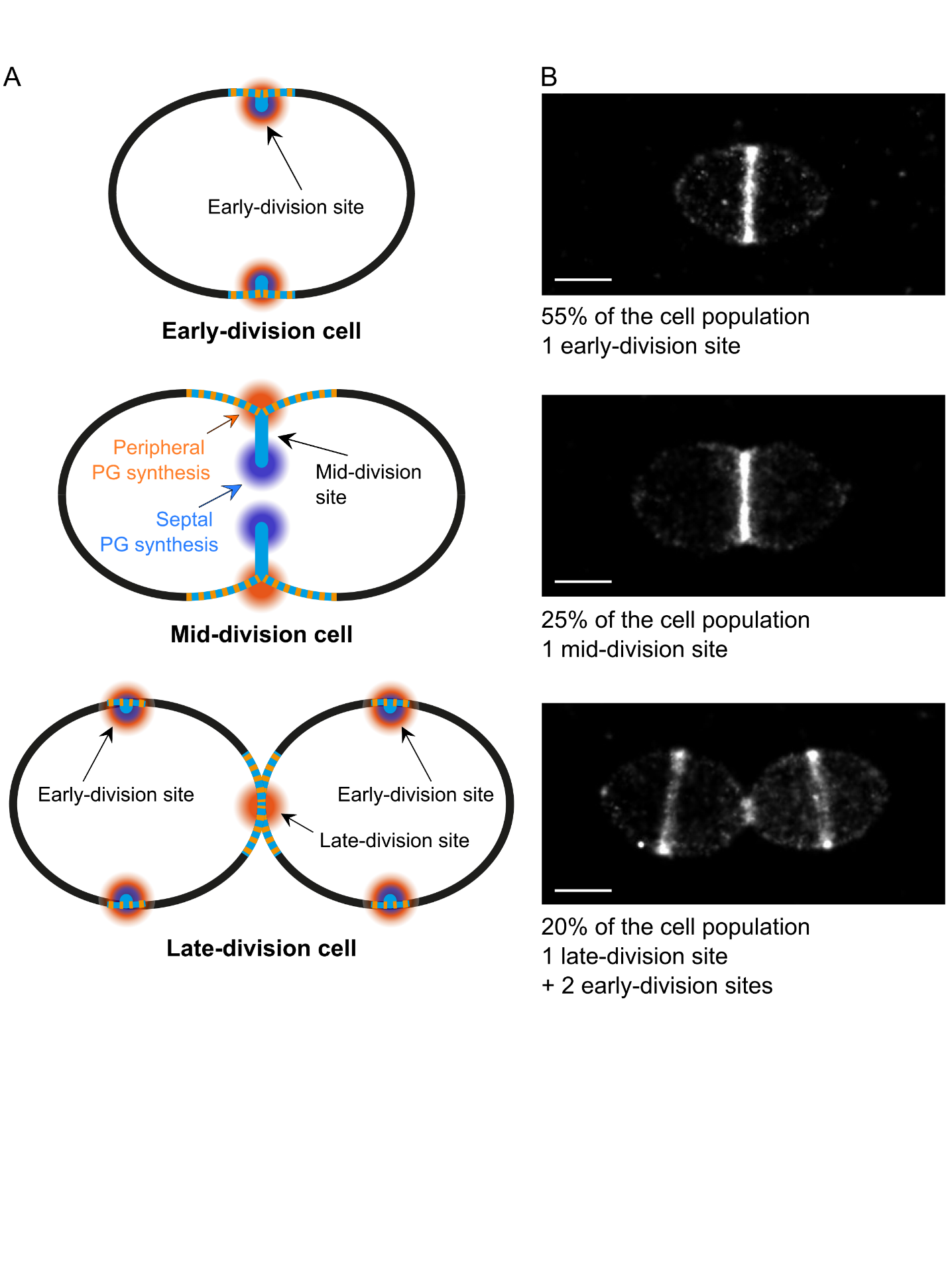

**Figure S1. Peptidoglycan synthesis in wild-type *S. pneumoniae.* A.** Model of PG growth and remodeling at three main stages of the cell cycle in ovococci. The septal PG (light blue) is synthesized by the divisome (dark blue pastille) at mid-cell, at the leading edge of the growing septum. The peripheral PG (light orange) is inserted by the elongasome (dark orange pastille) at the periphery of the septum, while this one is split and converted into lateral wall. At the beginning of the cell cycle (early-division cells), the divisome and elongasome co-localize within regions separated by less than 30 nm at early division sites. As septal PG is cleaved slower than it is synthesized, a septum forms progressively (mid-division cells), leading to the spatial separation of the septal and peripheral synthesis sites at mid-division sites. At the end of the cell cycle (late-division cells), the septum is closed but the elongasome continues synthesizing peripheral PG at the late-division site until full septum cleavage. At this stage, two new early PG labeling rings appear at the division sites of the future daughter cells. **B.** Representative dSTORM images of wild-type *S. pneumoniae* cells, corresponding to the cell cycle stages illustrated in panel A. The percentage of early, mid and late classes among the cell population (n = 231) is indicated, together with the number and category of division sites displayed by each class of cell. Scale bars, 500 nm.

.

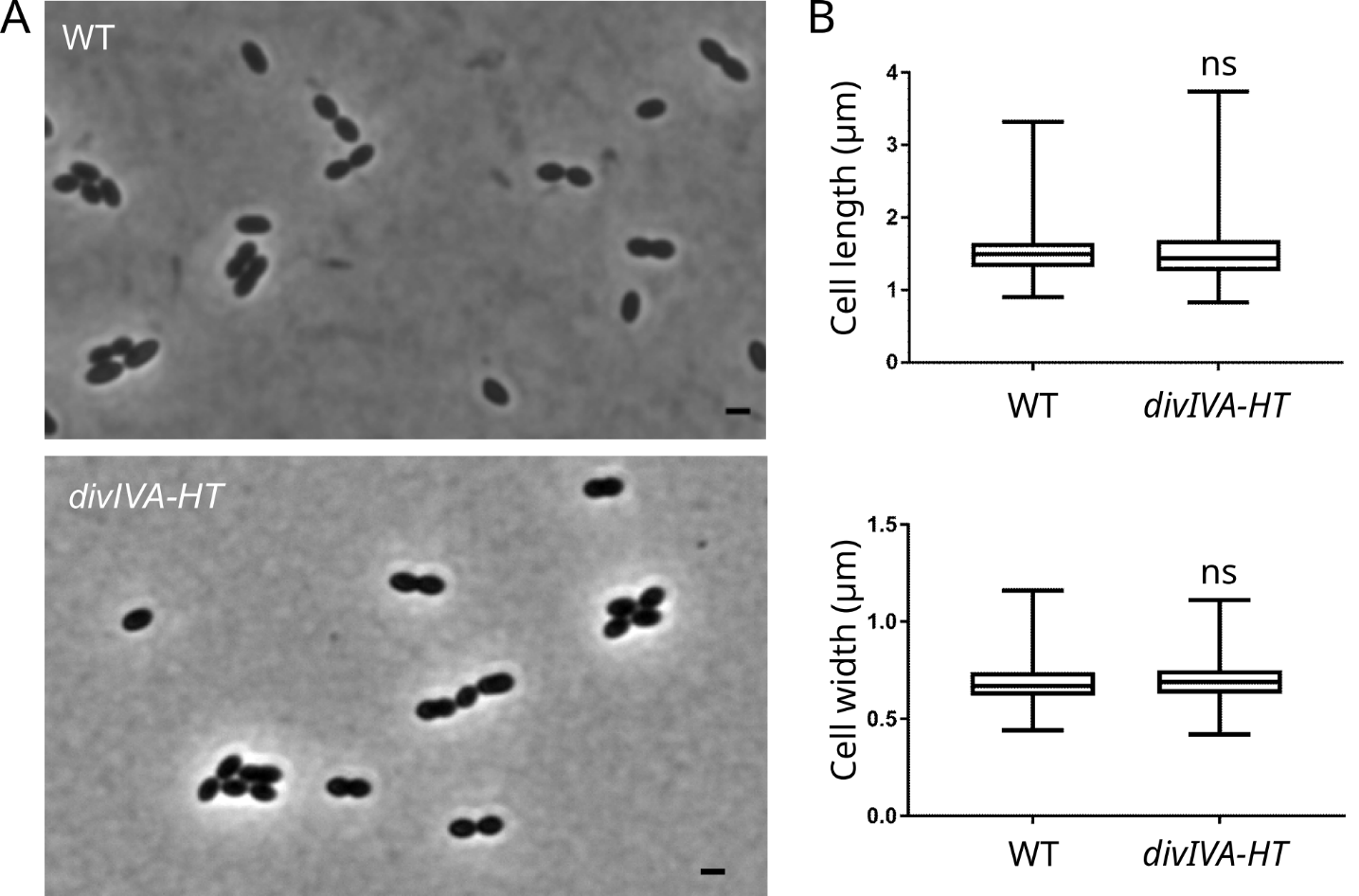
**Figure S2. Characterization of the cell morphology of the *S. pneumoniae* strain expressing the DivIVA-HT fusion protein. A.** Phase contrast images of exponentially growing wild-type (WT) *S. pneumoniae* and a strain producing a DivIVA-HT fusion protein from the endogenous *divIVA* site. Scale bars, 1 µm. **B.** Distributions of cell length and width among the cell population in WT (n = 620) and *divIVA-HT* (n = 717) strains are represented with box plots showing the interquartile range (25^th^ and 75^th^ percentile), the median value and whiskers for minimum and maximum values. P-values from the unpaired U test of Mann-Whitney show no significant difference in the cell length (p-value = 0.77) or in the cell width (p-value = 0.29) between the WT and *divIVA-HT* strains.

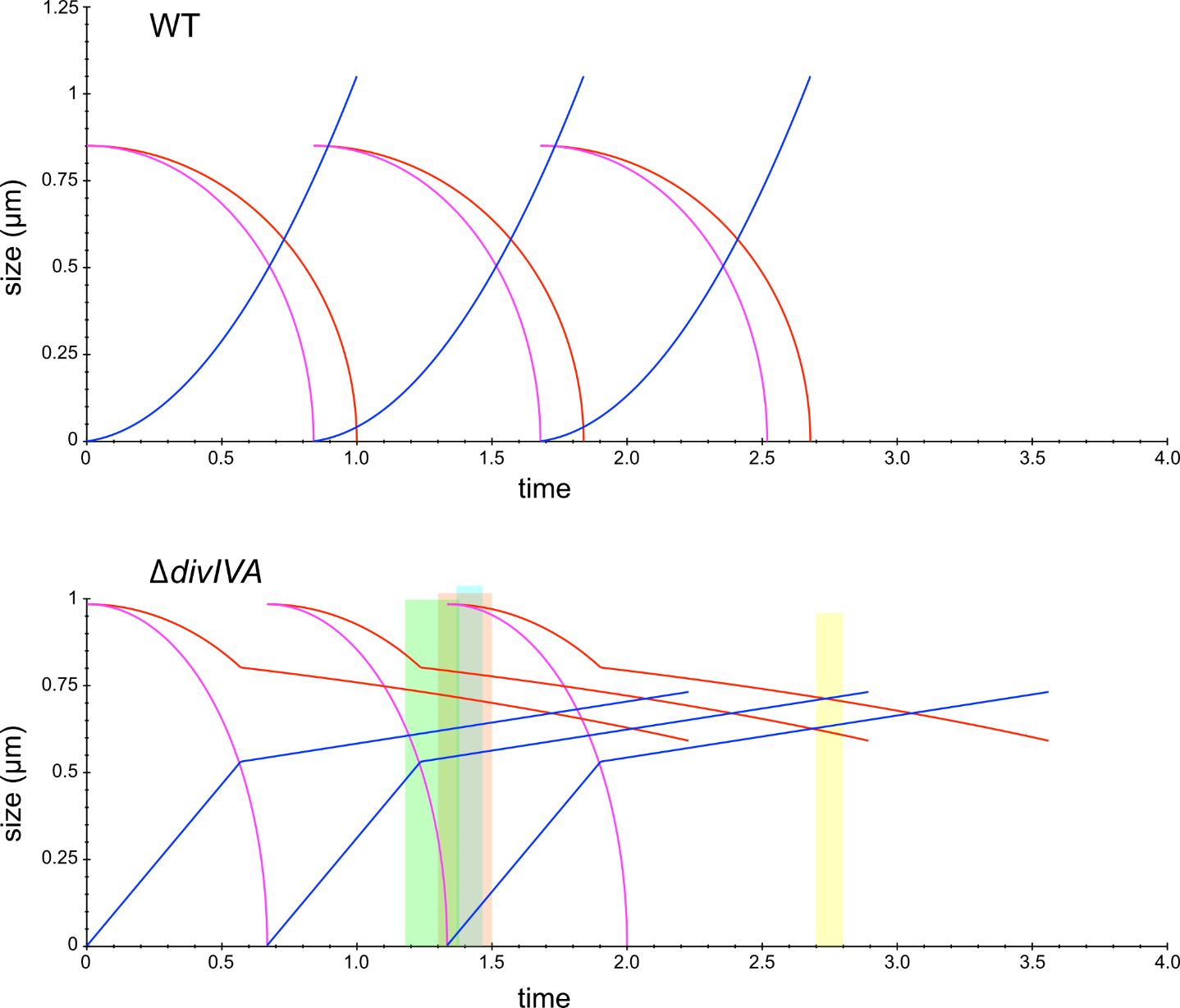
**Figure S3. Modeled evolution of the cell parameters of WT and Δ*divIVA* cells.** Blue, cell length *l*; red, outer septal diameter; magenta, inner septal diameter with the chosen parameters given in Table 1. The time unit is that required to complete a division for the WT strain, or that it would take Δ*divIVA* cells to complete a division if it were not slowed and stopped prematurely. The colored rectangles represent the timing of the labeling pulses of Fig. 3A (green), 3B (orange), 3C (cyan) and 3D (yellow).

**Supplementary Table 1. *S. pneumoniae* strains used in this study.**

| **Construct** | **Genotype** | | **Source** |
| --- | --- | --- | --- |
| R800 | | *rpsL1*; *Str^R^* | (Lefevre et al., 1979) |
| sspMJ83 | | R800, Δ*divIVA* markerless; *Str^R^* | (Fleurie et al., 2014) |
| sspJT90 | | R800, Δ*pbp2b mltG(Y488D)* markerless; *Str^R^* | This study |
| sspCM396 | | R800, Δ*pbp1a* markerless; *Str^R^* | This study |
| sspCM393 | | R800, Δ*pbp2a* markerless; *Str^R^* | This study |
| sspJT91 | | R800, Δ*divIVA* Δ*pbp2b mltG(Y488D)* markerless; *Str^R^* | This study |
| sspJT92 | | R800, Δ*divIVA* Δ*pbp1a* markerless; *Str^R^* | This study |
| sspJT96 | | R800, Δ*divIVA* Δ*pbp2a* markerless; *Str^R^* | This study |
| R1501 | | Δ*comC* | (Dagkessamanskaia et al., 2004) |
| R4596 | | Δ*comC*, *divIVA-HT* markerless | This study |

**Supplementary Table 2. Oligonucleotides used in this study.**

| **Primer name** | **Sequence** | | **Source** |
| --- | --- | --- | --- |
| DJ8 | | caatatcaccagacttcatgcaagac | This study |
| DJ10 | | accaaccacggtattggtcagaac | This study |
| DJ41 | | cagttggcccaatgtatgaagaaccagaagt caggatctggtggagaagcagc | This study |
| DJ42 | | gttggacctgtcggatgcactggagttaaccagagatttccaatgtagacaacc | This study |
| DJ43 | | ctgctgcttctccaccagatcctgacttctggttcttcatacattgggccaact | This study |
| DJ44 | | gttgtctacattggaaatctctggttaactccagtgcatccgacaggtccaac | This study |

**Supplementary Methods**

**Strain constructions**

Strain R4596, containing a *divIVA-HaloTag* fusion at the *divIVA* endogeneous locus, was generated as follows. Primers were designed to amplify PCR products containing: (I) the upstream region and the entire coding sequence of the *divIVA* (*spr1505*) gene (2234 bp; oligonucleotides DJ8 and DJ43, and R1501 DNA as template); (II) a 12 amino-acid linker, L5 (van Raaphorst et al., 2017), and the *orf* encoding the HaloTag sequence (982 bp; oligonucleotides DJ41 and DJ42, and a DNA fragment containing the HaloTag sequence generated by Integrated DNA Technologies); and (III) the downstream region of the *divIVA* gene (2062 bp; oligonucleotides DJ44 and DJ10, and R1501 DNA as template). The PCR products were gel-purified and used as templates in a SOEing PCR using the outer primers DJ8 and DJ10. The resulting SOEing PCR product was subsequently used to transform strain R1501 without selection, as previously described (Quevillon-Cheruel et al., 2012).
